# SOuLMuSiC: A Computational Tool for Predicting the Impact of Mutations on Protein Solubility

**DOI:** 10.1101/2025.01.15.633233

**Authors:** Simone Attanasio, Jean Kwasigroch, Marianne Rooman, Fabrizio Pucci

## Abstract

Protein solubility problems arise in a wide range of applications, from antibody development to enzyme production, and are linked to several major disorders, including cataracts and Alzheimer’s diseases. To assist scientists in designing proteins with improved solubility and better understand solubility-related diseases, we introduce SOuLMuSiC, a computational tool for the fast and accurate prediction of the impact of mutations on protein solubility. Our model is based on a simple shallow artificial neural network that takes as input a series of features, including biophysical properties of wild-type and mutated residues, energetic values computed using various statistical potentials, and mutational scores derived from protein language models. SOuLMuSiC has been trained on a curated dataset of about seven hundred mutations with known solubility values, collected and manually verified from original literature. It significantly outperforms current state-of-the-art predictors in strict cross-validation and shows good performance on external datasets containing high-throughput enzyme solubility-related data as well as protein aggregation propensities. In summary, SOuLMuSiC is a valuable tool for identifying mutations that impact protein solubility, and can play a major role in the rational design of proteins with improved solubility and in understanding genetic variants’ effect. It is freely available for academic use at http://babylone.ulb.ac.be/SoulMuSiC/.

## 1 Introduction

Solubility is a key biophysical property of proteins, the lack of which is often a major bottleneck for their production and storage [1, 2, 3]. Numerous applications in protein structure determination [1], pharmaceutics (e.g., production of protein-based therapeutics) [4, 5], and biotechnology (e.g., protein heterologous expression) [6, 7] require high-concentration protein formulation. Moreover, problems related to poor protein solubility and aggregation are central in a wide series of protein misfolding-related diseases, also known as proteinopathies. For example, Alzheimer’s and Parkinson’s diseases [8, 9] are characterized by the growth of insoluble deposits of misfolded proteins, cataracts are related to a decrease in the solubility of human *γ*-crystallin [10, 11], and amylin (Islet amyloid polypeptide - IAPP) is involved in diabetes [12]. The different solubility states of these proteins are also important for understanding how they spread and propagate between cells [13].

Despite the crucial importance of protein solubility and the considerable efforts made by the scientific community, a full comprehension of the mechanisms behind protein solubility remains out of reach. Solubility is influenced by a complex interplay of intrinsic factors such as residue-residue interactions, protein flexibility, amino acid composition, and hydrophobicity, as well as extrinsic variables such as pH, environmental temperature, ionic strength, and protein concentration [2, 14, 15, 16, 17].

In recent decades, multiple experimental and computational approaches have been developed to engineer proteins with improved solubility properties [3, 18, 19]. For example, when inclusion bodies form in heterologous expression, experimental protocols including the solubilization of these bodies through the addition of denaturant compounds followed by the refolding of the solubilized proteins have been designed to get bioactive recombinant proteins [20, 21]. Another experimental approach to enhance the solubility of a target protein involves fusing it with solubility-enhancing tags, even though this requires additional chromatographic steps to obtain tag-free recombinant proteins [22].

All these procedures remain, however, labor-intensive and quite unsatisfactory. Therefore, computational methods have been designed to speed up and complement the experimental approaches. A series of methods for predicting the solubility of a full protein have been developed, which leverage experimental solubility data [23] to train computational models [15, 24, 25, 26, 27, 28, 29, 30].

The challenge of predicting the impact of mutations on protein solubility has been underinvestigated due to the scarcity of mutagenesis data. Indeed, only recently has a database of mutations with known changes in solubility been collected [31]. Moreover, the variability in environmental conditions and in the methods used for variant characterization make constructing datasets and testing variant predictors challenging.

Over the past decade, a few methods have been developed to address this challenge, ranging from simple linear combinations of features to machine learning approaches, either considering the amino acid sequence or the three-dimensional structure. Renowned methods include CamSol [32], SODA [33] and PON-Sol2 [34]. However, these methods do not achieve satisfactory performance and moreover, the limited datasets they use for training make them prone to overfitting. A rigorous benchmark of their performances does not exist at the moment in the literature. For these reasons, we proceeded in this study to collect and manually curate a new dataset of mutations with an experimentally determined solubility value, and used it to develop a new method called SOuLMuSiC for the prediction of the impact of mutations on solubility. We compared SOuLMuSiC’s performance with the above mentioned algorithms and tested its ability to generalize to unseen data.

## 2 Methods

### Dataset curation

We collected a set of mutations, whose effects on protein solubility have been experimentally characterized. Although protein solubility can be rigorously defined in thermodynamics as the protein concentration in a saturated solution that is in equilibrium with a solid phase [2], its experimental measurement is far from trivial. Different techniques were used to estimate protein solubility, either directly or indirectly [35, 36]. They include measuring protein activity in a pellet after centrifugation [37], sodium dodecyl sulfate–polyacrylamide gel electrophoresis (SDS-PAGE) [38], and dynamic light scattering [39].

Due to the heterogeneity of experimental approaches, and the dependence of solubility on the environmental conditions used in the experiments, such as pH, temperature, and ion buffer, we chose not to report the exact values of the changes in solubility upon mutation:

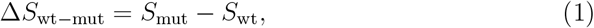

collected from the literature; besides, they are often not precisely known. Rather, we used five discrete solubility scores (−3, -1, 0, 1, 3) or equivalently, five solubility classes (- -, -, =, +, ++), and classified the effect of mutations in these classes according to their reported ΔSol values as defined in Table 1. Negative solubility scores represent a decrease in solubility upon mutation and positive values, a solubility increase. A value of zero indicates no significant change in solubility.

**Table 1:** Solubility scores and classes assigned to mutations. ΔSol represents the difference in solubility of the mutant protein relative to the wild type (in %).

| Effect on solubility | Solubility scores | Solubility classes |
| --- | --- | --- |
| $\Delta\text{Sol} \leq -50\%$ | -3 | - - |
| $-50\% < \Delta\text{Sol} \leq -10\%$ | -1 | - |
| $-10\% < \Delta\text{Sol} \leq +10\%$ | 0 | = |
| $+10\% < \Delta\text{Sol} \leq +50\%$ | 1 | + |
| $\Delta\text{Sol} > +50\%$ | 3 | ++ |

To set up our first mutation dataset 𝒟_*Sol*_, we started to collect mutations from the mutational solubility database SoluProtMutDB [31]. We then performed an additional literature search involving manually reviewing each SoluProtMutDB entry in the original literature to correct errors, and normalizing the solubility values according to our 5-value classification. Additionally, we extended this search to incorporate recent literature and patent data that were not included in SoluProtMutDB. We excluded data originating from high-throughput techniques such as fluorescence-activated cell sorting, and from mutations in membrane proteins. We focused exclusively on single-site mutations, thus not recording any multiple-site mutations.

We collected the 3-dimensional (3D) structures of all wild-type proteins in 𝒟_*Sol*_. We considered the experimental structures in the Protein Data Bank (PDB) [40] when available, but only if their resolution was smaller than 2.5 Å and their sequence 100% identical to the one considered. Otherwise, we modeled the structure using AlphaFold 2 [41]. In the few cases in which the oligomeric structures were too large to be modeled with AlphaFold, we used the homology modeling algorithm SWISS-MODEL [42]. The final dataset 𝒟_*Sol*_ consists of 702 mutations with experimentally characterized effects on protein solubility, inserted in 80 proteins with experimental or modeled structures.

We set up a second dataset, called 𝒟_*Inv*_, containing the reverse mutations of 𝒟_*Sol*_. More precisely, for each mutation (wt → mut) in 𝒟_*Sol*_ with a given ΔSol value, we included the reverse mutation (mut → wt) in 𝒟_*Inv*_ and assigned it a value of −ΔSol. We created this database to test the symmetry properties of our predictor, as non-symmetric models that are biased towards the training set are often constructed, as shown in [43]. The structures of the 702 mutant proteins in 𝒟_*Inv*_ were modeled with the homology algorithm Modeller [44] using as template the structures contained in 𝒟_*Sol*_.

As additional test mutation dataset, called 𝒟_*LGK*_, we used solubility data of Levoglucosan kinase (LGK) from *Lipomyces starkeyi*. This enzyme catalyzes the phosphorylation of levoglucosan [45]. Deep mutational scanning was performed on LGK using yeast surface display (YSD) [46]: proteins were fused with an N-terminal domain to localize the protein on the outer cell surface, and with a C-terminal epitope tag, which binds to a fluorescent antibody, enabling the identification of only the variants expressed on the cell surface. Misfolded protein variants are less expressed on the cell surface as the proteasome degrades them to ensure protein quality control. In total, 𝒟_*LGK*_ consists of 6,246 single-site mutations with known solubility values. Note that these values are only partially related to real solubility as the proteasome can also degrade misfolded soluble proteins. The structure that we used for LGK is homodimeric and has the PDB code 4ZFV.

In the last database, referred to as 𝒟_*Aβ*_, we collected the nucleation scores of variants in A*β*42, a protein known to play a key role in the pathogenesis of Alzheimer’s disease. The variant scores were measured [47] through a cell-based selection assay in which A*β* aggregation triggers the aggregation of an endogenous protein, a process that is crucial for cell growth under selective conditions. In total, we collected 632 variants with known changes in nucleation scores. Note that these scores are only partially related to solubility, as aggregation phenomena differ from poor solubility precipitation. The structure that we used for A*β*42 has the PDB code 1IYT.

The datasets 𝒟_*Sol*_, 𝒟_*Inv*_, 𝒟_*LGK*_ and 𝒟_*Aβ*_, as well as all structures used in this study, can be retrieved from our GitHub repository at github.com/3BioCompBio/SOuLMuSiC.

### Features

In our SOuLMuSiC model, we used two types of features: structure-based and sequence-based. The former are mainly based on statistical potentials, and the latter, on various amino acid scales and on protein large language models. Below, we provide a brief description of these features.

#### Statistical potentials

which are well-known mean force potentials extracted from frequencies of associations between sequence elements, *se*, and structure motifs, *st*, in datasets of protein structures [48, 49]. Using this formalism, the folding free energy Δ*W* (*st, se*) of the sequence-structure pair (*st, se*) can be computed in terms of the probabilities of *se, st* and (*st, se*) using the inverse Boltzmann law, as defined in the first equality :

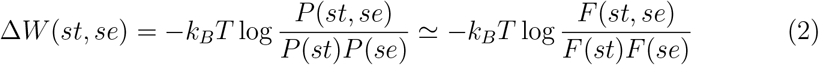

where *k*_*B*_ is the Boltzmann constant and *T* the absolute temperature taken to be room temperature [49]. As shown in the second approximate equality, the probabilities *P* can be estimated from the frequencies *F* of observation of *se, st* and (*st, se*) in a high-quality, non-redundant dataset of experimental protein 3D structures.

In our model, we used four types of statistical potentials, differing in the structure and sequence elements considered. Their list and characteristics are given in Table 2. Among these four potentials, two describe local interactions along the polypeptide chain and two are distance potentials that describe tertiary interactions and the likelihood of amino acids being separated by specific spatial distances. Note that more potentials can be defined, but we combined some of them to limit the number of free parameters that we have to optimize and thus to avoid overfitting.

**Table 2:**
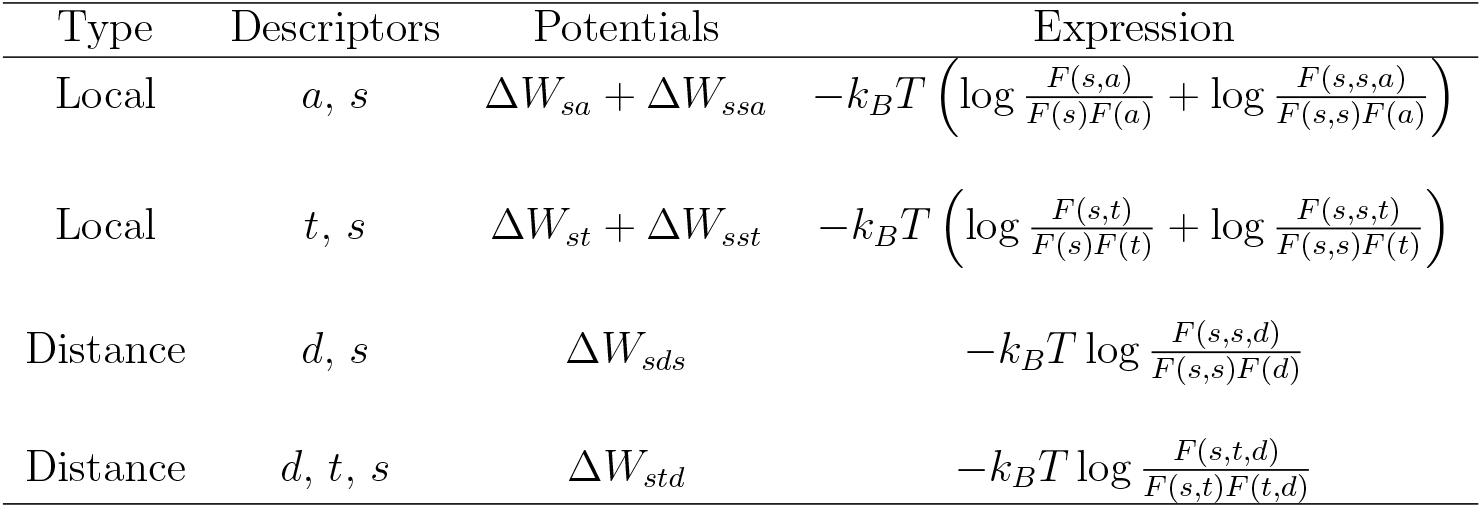
List of statistical potentials used as features in SOuLMuSiC. The sequence descriptor used is the amino-acid type *s*, and the sequence elements, *se*, are single amino acids *s* or amino acid pairs (*s, s*). The structure descriptors are: the solvent accessibility *a* of a residue defined as the ratio of its solvent accessible surface area in a given structure and in an extended Gly-X-Gly tripeptide conformation, computed as in [52]; the backbone torsion angle domain *t* as defined in [53]; the distance *d* between the geometric center of the heavy side chain atoms of two residues as defined in [54]. The structure motifs, *st*, considered are *a, t, d*, and (*t, d*). More details can be found in [50, 51]

| Type | Descriptors | Potentials | Expression |
| --- | --- | --- | --- |
| Local | $a, s$ | $\Delta W_{sa} + \Delta W_{ssa}$ | $-k_B T \left( \log \frac{F(s,a)}{F(s)F(a)} + \log \frac{F(s,s,a)}{F(s,s)F(a)} \right)$ |
| Local | $t, s$ | $\Delta W_{st} + \Delta W_{sst}$ | $-k_B T \left( \log \frac{F(s,t)}{F(s)F(t)} + \log \frac{F(s,s,t)}{F(s,s)F(t)} \right)$ |
| Distance | $d, s$ | $\Delta W_{sds}$ | $-k_B T \log \frac{F(s,s,d)}{F(s,s)F(d)}$ |
| Distance | $d, t, s$ | $\Delta W_{std}$ | $-k_B T \log \frac{F(s,t,d)}{F(s,t)F(t,d)}$ |

Once these potentials were derived, we used them to compute the change in folding free energy ΔΔ*G* between the wild type (wt) and mutant (mut) proteins. Considering for example the distance potentials Δ*W*_*sds*_ defined in Table 2, the associated free energy change ΔΔ*G*_*SDS*_ is defined as:

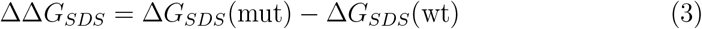

where

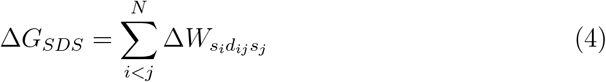

with *N* the protein length, *s*_*i*_ and *s*_*j*_ the sequence types of residues *i* and *j*, and *d*_*ij*_ the distance between the geometric side chain centers of residues *i* and *j*. The sum is thus over all residue pairs in the protein. Analogous expressions can be derived for the other potentials.

For technical details on the construction and implementation of statistical potentials we refer the reader to [50, 51].

#### Amino acid scale-based features

We used four sequence-based features that are essentially amino acid scales, and a fifth one corresponding to the large language model (LLM) ESM-1v. They are described below; the values for the four former scales can be found in our GitHub repository github.com/3BioCompBio/SOuLMuSiC.

⋄ ΔHydro is the difference between the hydrophobicity of the mutant and wild-type residues. The hydrophobicity scale we used is taken from [55]. It shows the best correlation with experimental solubility values, compared to the other tested scales such as the Kyte-Doolittle scale [56] and the Janin scale [57],
⋄ ΔAro is the difference in aromaticity between the mutant and wild-type residues. The aromaticity is considered equal to one for PHE, TYR, and TRP, and equal to zero for all other amino acids.
⋄ ΔIso is related to the isoelectric point Iso of the amino acid under consideration. As the solubility of proteins is minimal near the isoelectric pH [58, 59] and our predictions are for neutral pH, i.e. seven, we defined the feature ΔIso as the difference of (Iso−7)^2^ for mutant and wild-type proteins:

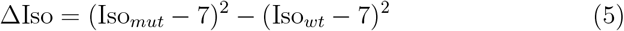
⋄ ΔΔApaac is defined from the amphiphilic pseudo amino acid composition. Apaac describes the hydrophobicity and hydrophilicity distribution patterns along the protein chain and therefore includes sequence-order effects [60]. For a given protein, the Apaac per residue type is its frequency normalized by a factor that includes the correlation effects between hydrophobic or hydrophilic residues along the chain. We downloaded the dataset [15], in which about 11 thousand protein sequences were experimentally identified as soluble or insoluble. For each of these two sets, we calculated the Apaac score per amino acid type using the protr package [61] and averaged them over all sequences in each of the two sets. The logarithm of the ratio between Apaac scores of soluble and insoluble sets yields the ΔApaac index. ΔΔApaac is then computed as the difference between the mutant and wild type ΔApaac values.

#### Protein Large Language models

As the last sequence-based feature, we leveraged ESM (ESM-1v), a freely available unsupervised LLM machine learning model that predicts the variant effects from the amino acidic sequence. It is based on a 650M parameter transformer language model with zero-shot inference [62] trained on Uniref90 2020-03 [63].

### Artificial Neural Network models

We have a total of nine features that we decided to combine using a simple shallow artificial neural network. We used an approach analogous to that of the PoPMuSiC and HoTMuSiC algorithms [50, 64], i.e., we defined the perceptron activation functions to be sigmoid functions of the solvent accessibility of the mutated residues. This involves weighting the input features differently in terms of the per-residue solvent accessibility, as different potentials and features of amino acids are known to contribute differently in the protein core and on the surface [65].

More precisely, the SOuLMuSiC model reads as:

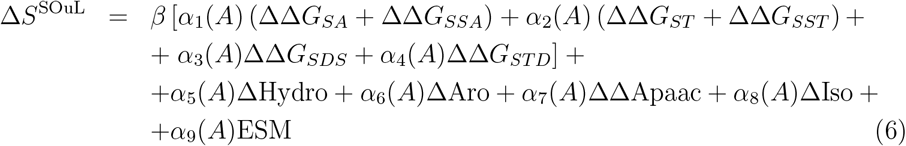

where *β* can be equal to 1 or −1, depending on whether the input structure is a globular protein or an insoluble supramolecular homopolymer of proteins, such as amyloid fibrils, and *α*_*i*_(*A*) (*i* = 1, .., 9) are sigmoid functions expressed as :

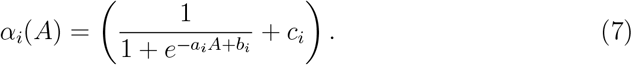

This yields 27 free parameters (*a*_*i*_, *b*_*i*_, *c*_*i*_), which were identified by minimizing the root mean square error (RMSE) between experimental and predicted changes in solubility Δ*S* upon mutations for the mutation dataset 𝒟_*Sol*_ used as training set:

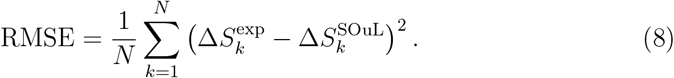

where *N* is the number of mutations in 𝒟_*Sol*_. Minimization is performed using the standard gradient descent algorithm implemented in [66] with default parameters. Performance is evaluated using a strict leave-one-out cross-validation process, where all mutations in one protein are, in turn, excluded from the training set and then predicted.

## 3 Results

### Prediction performances

We tested the prediction performance of SOuLMuSiC on different datasets. We started with the training set 𝒟_*Sol*_, and evaluated the performance in leave-one-out cross-validation at protein level. The distributions of the cross-validated SOuLMuSiC scores for all 𝒟_*Sol*_ entries, separated as a function of their experimentally characterized solubility score defined in Table 1, are shown in Fig. 1. We clearly observe that SOuLMuSiC can distinguish between different solubility classes and identifies with fair accuracy which mutations make proteins more soluble or less soluble.

**Figure 1:**
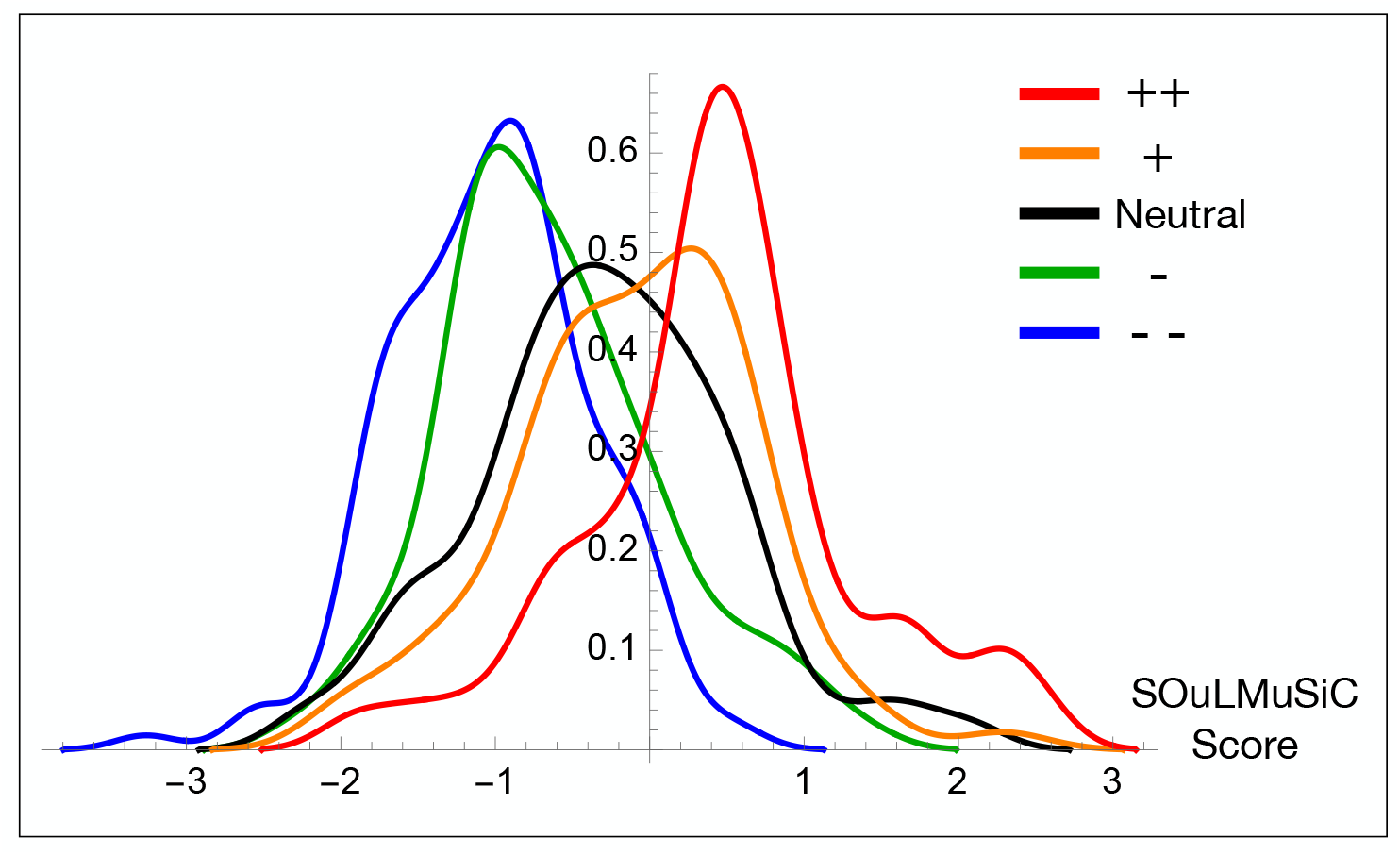
SOuLMuSiC’s score distribution in the 𝒟_*Sol*_ dataset, obtained in protein-level leave-one-out cross validation, for the five classes of mutations experimentally shown to affect solubility from strongly increasing (++) to strongly decreasing (--), as defined in Table 1.

One of the metrics that we used to evaluate the overall performance of SOuLMuSiC is the Spearman correlation coefficient *ρ* between the predicted and experimental Δ*S* values. SOuLMuSiC achieves a *ρ* value of 0.46 in the dataset 𝒟_*Sol*_ with the above-mentioned strict cross-validation procedure. This is a fairly strong correlation considering the variability in experimental setups used to determine the ΔSol values.

We also used other performance metrics to evaluate SOuLMuSiC, i.e. multiclass precision, recall (or sensitivity), and balanced accuracy (BACC). They are defined and given in Table 3. To convert the SOuLMuSiC score into a multiclass prediction, we set thresholds at the midpoint between the mean SOuLMuSiC scores of each pair of adjacent classes. The global value of each metric is obtained by averaging the metric value over each of the five classes.

**Table 3:** Cross-validated five-class prediction accuracy of SOuLMuSiC in the 𝒟_*Sol*_ dataset. The five classes are (++, +, =, -, - -), as defined in Table 1. Precision is defined as (TP/(TP+FP)) where TP, TN, FP, FN are true positives, true negatives, false positive and false negatives, respectively. Recall is defined as (TP/(TP+FN)) and balanced accuracy (BACC), as (precision + recall)/2.

| Classes | Precision | Recall | BACC |
| --- | --- | --- | --- |
| ++ | 0.24 | 0.69 | 0.47 |
| + | 0.14 | 0.16 | 0.15 |
| = | 0.48 | 0.15 | 0.31 |
| - | 0.25 | 0.18 | 0.21 |
| -- | 0.36 | 0.64 | 0.50 |
| Global | 0.29 | 0.36 | 0.33 |

**Table 4:** Spearman correlation coefficients *ρ* between experimental Δ*S* values and those predicted by different predictors on the 𝒟_*Sol*_ dataset. Note that SOuLMuSiC scores are computed in leave-one-out cross validation on 𝒟_*Sol*_, PON-Sol2 is trained on a dataset that largely overlap with it, while the others are essentially blind to it. “Res” means features derived from biophysical data at amino acid level (e.g., hydrophobicity, isoelectric point); “Seq” means sequence-based features (e.g., residue secondary structure propensity, residue flexibility); “Str” means structural features; “Evo” means evolutionary features; ΔΔG means changes in folding free energy upon mutations. Here we only report the results of the predictors tested that show a statistically significant correlation with experimental data (P-value *<* 0.01).

| Method | Prediction Types | Feature Types | $\rho$ |
| --- | --- | --- | --- |
| SOuLMuSiC | $\Delta S$ | Str, NLP, Res | 0.46 |
| CamSol [32] | $\Delta S$ | Str, Res, Seq | 0.27 |
| PON-Sol2 [34] | $\Delta S$ | Evo, NLP, Res, Seq | 0.25 |
| Protein-Sol [26] | S | Res, Seq | 0.16 |
| SoDoPE [15] | S | Res | 0.13 |
| SoluProt [28] | S | Res, Seq | 0.10 |
| ESM-1v [62] | Fitness | NLP | 0.30 |
| PoPMuSiC [50] | $\Delta\Delta G$ | Str | 0.30 |

The global BACC is equal to 0.33, which can be compared to the expected BACC of 0.2 for a random prediction by a five-class classifier. Although this score is not outstanding, a closer look at the data reveals that the classes of variants with a significant impact on solubility are well predicted, with BACC scores of about 0.5. Other classes are, however, less accurately predicted, which may be partly due to the way we defined these classes: for example, we label a change in solubility as “neutral” if it ranges between -10% and +10% compared to the wild type, while the “+” and “-” classes cover changes between +10% to +50% and -10% to -50%, respectively. Given that solubility values can vary greatly depending on experimental methods and environmental conditions, it is unsurprising that the model struggles more with distinguishing classes with small changes in solubility values, as some misclassifications may also arise from inherent variability in the dataset.

The global recall shows very similar trends to the global BACC score, with classes “++” and “- -” being largely better predicted. The neutral class has much better precision and much worse recall, meaning that the number of false negatives is much larger than the number of false positives. This is due to the choice of the class threshold, which tends to favor the prediction of the extreme classes over the neutral one, thereby narrowing the interval in which predictions are classified as neutral.

### Comparison with other predictors

We compared SOuLMuSiC’s performances on the 𝒟_*Sol*_ dataset with state-of-the-art solubility predictors. Most of these methods use biophysical data at amino acid level such as hydrophobicity or isoelectric point, and sequence-based information such as residue propensities to adopt secondary structures or residue flexibility in a given sequence window. Only few use structural information (i.e., SOuLMuSiC, CamSol [32], SOLart [27]). Recent tools, such as SOuLMuSiC, NetSolP [30], and PON-Sol2 [34], also integrate natural language-based predictions (NLP), following recent advances in the protein NLP field [62].

Among these prediction methods, only 4 (SOuLMuSiC, CamSol [32], PON-Sol2 [34], SODA [33]) directly predict solubility changes upon mutations, with the former three providing a continuous score and the latter being a binary classifier. Seven others (Protein-Sol [26], SoDoPE [15], SoluProt [28], ccSOLomics [25], NetSolP [30], Skade [29], SOLart [27]) predict protein solubility values; the ΔS values in these cases are obtained by subtracting the S values of wild-type and mutant. Two predictors, PoPMuSiC and ESM, predict the folding free energy changes upon mutation (ΔΔ*G*s) and the fitness changes upon mutation, respectively.

We computed the Spearman correlation coefficients *ρ* between the Δ*S* predicted by the methods tested and the experimental Δ*S* for all the entries in the 𝒟_*Sol*_ dataset. We see that SOuLMuSiC largely outperforms the other predictors, achieving a solid Spearman correlation coefficient of 0.46 in leave-one-out cross-validation. Note that PON-Sol2 is trained on a dataset that largely overlap with 𝒟_*Sol*_, while all other predictors are essentially blind to it. Although predictors claim higher accuracy in their papers, when these models are tested on new, larger, and highly curated datasets such as the 𝒟_*Sol*_ dataset, their performance often drops significantly. It must be noted that the number of entries in solubility datasets remains limited (approximately 700 for 𝒟_*Sol*_ and only a few hundreds for other datasets used in the literature to train models), which hinders the development of sufficiently robust methods and limits the possibility to avoid overfitting. Moreover, most methods do not predict the change in solubility but rather the solubility of the input sequence, a slightly different problem that makes predicting changes in solubility more challenging for them. Indeed, almost half of the predictors mentioned above do not show a statistically significant correlation with 𝒟_*Sol*_ values and are thus not reported in Table 3. The last two methods tested, ESM-1v and PoPMuSiC, predict changes in fitness and thermodynamic stability upon mutations, respectively, both of which are only partly related to solubility. Despite this, their correlation of 0.3 is surprisingly the highest after SOuLMuSiC’s correlation, indicating a close relationship of their descriptors with solubility.

### Test on inverse mutations and symmetry properties

We evaluated SOuLMuSiC on the dataset of reverse mutations, 𝒟_Inv_, to test the symmetry properties of our model. If the change in protein solubility due to a mutation from protein ”wt” to protein ”mut” is given by Eq. 1, the reverse mutation from protein ”mut” to protein ”wt” should, by construction, satisfy

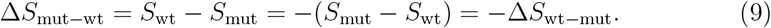

This symmetry property has already been thoroughly discussed and investigated in the field of predicting changes in protein stability and protein-protein binding affinity upon mutations [43, 67, 68].

To quantify how much the predictions deviate from perfect symmetric behavior, we introduced two scores [43]. The first, *r*_dir,inv_, is the Pearson correlation coefficient between the predictions on the datasets of direct and inverse mutations, which is equal to −1 for a perfectly symmetric predictor. The second score, ⟨*δ* ⟩, is defined as:

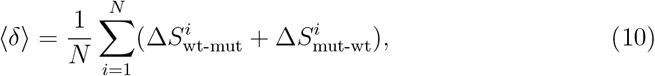

where the sum is over all N mutations in a given dataset. For a perfectly symmetric predictor, ⟨*δ* ⟩ is equal to 0.

We evaluated SOuLMuSiC and its features on the datasets of direct and inverse mutations, 𝒟_Sol_ and 𝒟_Inv_. Table 5 provides the values of the two metrics ⟨*δ*⟩ and *r*_dir,inv_, along with *ρ*_dir_ and *ρ*_inv_, the Spearman correlation coefficients between predicted and experimental values on these two datasets. SOuLMuSiC shows rather good performance in terms of symmetry with *r*_*dir,inv*_ = −0.81 and ⟨*δ* ⟩= −0.35. Indeed, the latter value is rather are small compared to the standard deviation of the predicted score distributions, which is in the 0.8-0.9 range for 𝒟_Sol_ and 𝒟_Inv_.

**Table 5:** Symmetry properties of SOuLMuSiC and of the features used for SOuLMuSiC’s construction, measured by *r*_*dir,inv*_ and ⟨*δ*⟩; SOuLMuSiC’s performance, evaluated by *ρ*_*dir*_ and *ρ*_*inv*_.

| | $r_{dir,inv}$ | $\langle\delta\rangle$ | $\rho_{dir}$ | $\rho_{inv}$ |
| --- | --- | --- | --- | --- |
| SOuLMuSiC | -0.81 | -0.35 | 0.53 | 0.45 |
| **Amino acid scales** |  |  |  |  |
| $\Delta\text{Hydro}$ | -1.00 | 0.00 | -0.18 | -0.18 |
| $\Delta\text{Aro}$ | -1.00 | 0.00 | -0.18 | -0.18 |
| $\Delta\text{Iso}$ | -1.00 | 0.00 | 0.12 | 0.12 |
| $\Delta\Delta\text{Apaac}$ | -1.00 | 0.00 | 0.19 | 0.19 |
| **Statistical potentials** |  |  |  |  |
| $\Delta W_{st} + \Delta W_{sst}$ | -0.81 | 0.03 | -0.25 | -0.22 |
| $\Delta W_{sa} + \Delta W_{ssa}$ | -0.83 | 0.03 | -0.31 | -0.31 |
| $\Delta W_{sds}$ | -0.83 | 0.01 | -0.28 | -0.27 |
| $\Delta W_{std}$ | -0.72 | -0.03 | 0.12 | 0.10 |
| **pLM** |  |  |  |  |
| ESM-1v | -0.93 | -6.4 | 0.30 | 0.27 |

The Spearman correlation coefficients between the SOuLMuSiC predictions and the experimental values for 𝒟_Sol_ and 𝒟_Inv_ are also good, although somewhat better for the direct mutation set than for the inverse mutation set, with values of 0.53 and 0.45, respectively.

To better understand where the deviation from a symmetric behavior comes from, we analyzed the properties of each of the nine components of SOuLMuSiC. The features defined from amino acid scales are perfectly symmetric but correlate poorly with experimental data (*ρ*_*dir*_ and *ρ*_*inv*_ between 0.1 and 0.2). The statistical potentials are almost symmetric in terms of ⟨*δ*⟩. They are, however, not totally symmetric in terms of *r*_inv,dir_, because they incorporate some inaccuracies as mutated structures are modeled rather than experimental, and intrinsic noise caused by statistical errors in their derivation. In contrast, the pLM feature has good symmetry properties in terms of *r*_inv,dir_, but not at all in terms of ⟨*δ*⟩. It is biased toward negative values, which means that neutral mutations are not defined by a score of 0. Both the statistical potentials and pLM correlate significantly better with experimental data than amino acid scales on both direct and inverse mutations. Indeed, *ρ*_*dir*_ and *ρ*_*inv*_ are between 0.2 and 0.3 for all statistical potentials except the “std” potential, and are equal to 0.3 for pLM.

### Application to LGK and trade-off between solubility and stability

As an additional test of SOuLMuSiC, we studied the solubility of Levoglucosan kinase (LGK) from *Lipomyces starkeyi*, an enzyme that catalyzes the phosphorylation of levoglucosan [45]. In more detail, we analyzed the interplay between protein solubility and stability, which generally exhibits a trade-off [69]. In general, increasing thermodynamic stability of proteins tends to reduce their aggregation propensities and make them more soluble, but there are examples of proteins with enhanced stability and reduced solubility [70].

To analyze this trade-off, we applied our SOuLMuSiC predictor, along with PoPMuSiC [50], a tool we previously developed to predict the impact of mutations on thermodynamic stability. We compared these predictions with deep mutagenesis scanning data where the effects of amino acid substitutions on a combination of protein solubility and stability were systematically explored using yeast surface display (YSD) [46]. Solubility and stability are challenging to disentangle in high-throughput data, and that is why the YSD experiment considered measures a combination of both.

The results of the comparison are shown in Table 6. We first compared the results of the SOuLMuSiC and PoPMuSiC scores with the experimental YSD scores [46], finding a good Spearman correlation coefficient of *ρ* = 0.35 and *ρ* = −0.29, respectively. Note that solvent accessibility has a good Spearman correlation of 0.30, indicating the tendency of mutations in the protein core to have larger impact than those at the surface. Note that this is even more marked for stability [71] than for solubility.

**Table 6:** Spearman correlation coefficients *ρ* between the results of the YSD score on the LGK dataset [46] and SOuLMuSiC’s and PoPMuSiC’s predictions, solvent accessibility and the combination of SOuLMuSiC and PoPMuSiC (SOuLPoP) according to Eq. 11.

|  | YSD score |
| --- | --- |
| SOuLMuSiC | 0.35 |
| PoPMuSiC | -0.29 |
| Solvent accessibility | 0.30 |
| SOuLPoP | 0.40 |

Since the experimental YSD score reflects a mixture of the two quantities, we also combined our prediction scores in the following way:

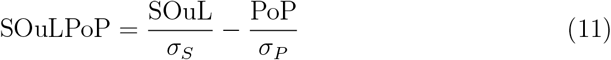

where PoP and SOuL are the PoPMuSiC ΔΔ*G* prediction and SOuLMuSiC score, respectively, and *σ*_*P*_ and *σ*_*S*_ their respective root mean square deviations on the LGK dataset. Note the minus sign between the two terms. Variants that increase stability (with a negative ΔΔ*G* in our conventions) are likely to also increase solubility, resulting in a positive SOuL score. As the YSD score arises from a combination of solubility and stability, we observe that the combined SOuLMuSiC and PoPMuSiC predictions perform better than each individually, reaching the correlation *ρ* = 0.40.

Note that solubility and stability are mildly correlated, as discussed previously.The Spearman correlation coefficient between the SOuL scores and PoPMuSiC’s ΔΔ*G* values on the LGK dataset is equal to *ρ* = −0.3.

It is well known that mutations near the catalytic site are likely to disrupt enzyme activity but are often stabilizing [72]. To study the catalytic site in more detail, we analyzed the prediction scores for all possible mutations of residues located within a distance of less than 6 Å from the center of the catalytic site and compared them with those further away. We found that mutations in the catalytic site have an average ΔΔ*G* value of 0.47 kcal/mol, which is lower than the 1.20 kcal/mol observed for other residues, suggesting that these regions are inherently less impacted in terms of stability [72]. For solubility, however, there is essentially no difference, with the average score for residues close to and far from the catalytic site being equal to about -1.0.

### Application to aggregation-prone proteins

Protein aggregation is a well-studied yet poorly understood phenomenon that leads to a series of pathological conditions [73, 74]. A series of proteins end up in mis-folded states that can trigger the formation of supramolecular assemblies such as amyloid fibrils. These insoluble assemblies are often observed as deposits in major neurodegenerative diseases: for example, the aggregation of *α*-synuclein is considered as one of the hallmarks of Parkinson’s disease, the formation of amyloid beta (A*β*) plaques and neurofibrillary tangles composed of the Tau protein is strongly involved in the pathogenesis of Alzheimer’s disease, and Huntingtin fibrils are the toxic species involved in Huntington’s disease. Although amyloid formation and protein solubility are distinct phenomena, they are interrelated [32, 75]. Thus, we evaluated our SOuLMuSiC predictor on its ability to predict the effects of variants on aggregation-prone proteins.

Aggregation-prone proteins are often characterized by flexible and intrinsically disordered regions, making it difficult to obtain high-quality structures of their folded form [76]. However, obtaining the structure of amyloid fibrils is comparatively easier and indeed, hundreds of them can be found in the PDB.

To study protein aggregation propensities, we applied SOuLMuSiC to the A*β*-42 protein, using both the amyloid fibril structure and the protein-in-solution structure as input. When the input structure is an insoluble aggregate, we set the parameter *β* in front of the structural terms in our model to *β* = −1 (see Eq. 6). In contrast, SOuLMuSiC applied to folded proteins in solution uses *β* = 1. This adjustment is thermodynamically intuitive: stabilizing the native globular form of a protein generally leads to increased solubility, while stabilizing an insoluble form, such as a fibril, further decreases the solubility of the assembly.

We applied SOuLMuSiC to the structure of the A*β*-42 fibrils identified with the PDB code 2NAO [77] and to the structure of the in-solution A*β*-42 protein identified with the code 1IYT [78]. Note that both the 2NAO and 1IYT structures were derived from NMR data; we used as SOuLMuSiC scores the average of the scores on all 10 NMR structures in the PDB files. We then compared these scores with the solubility data [79] derived from a yeast aggregation assay combined with deep mutational scanning. We focused on single amino acid substitutions, excluding truncating and synonymous mutations, which amounts to a total of 790 mutations. We potted the predicted scores versus the experimental values in Figure 2.

**Figure 2:**
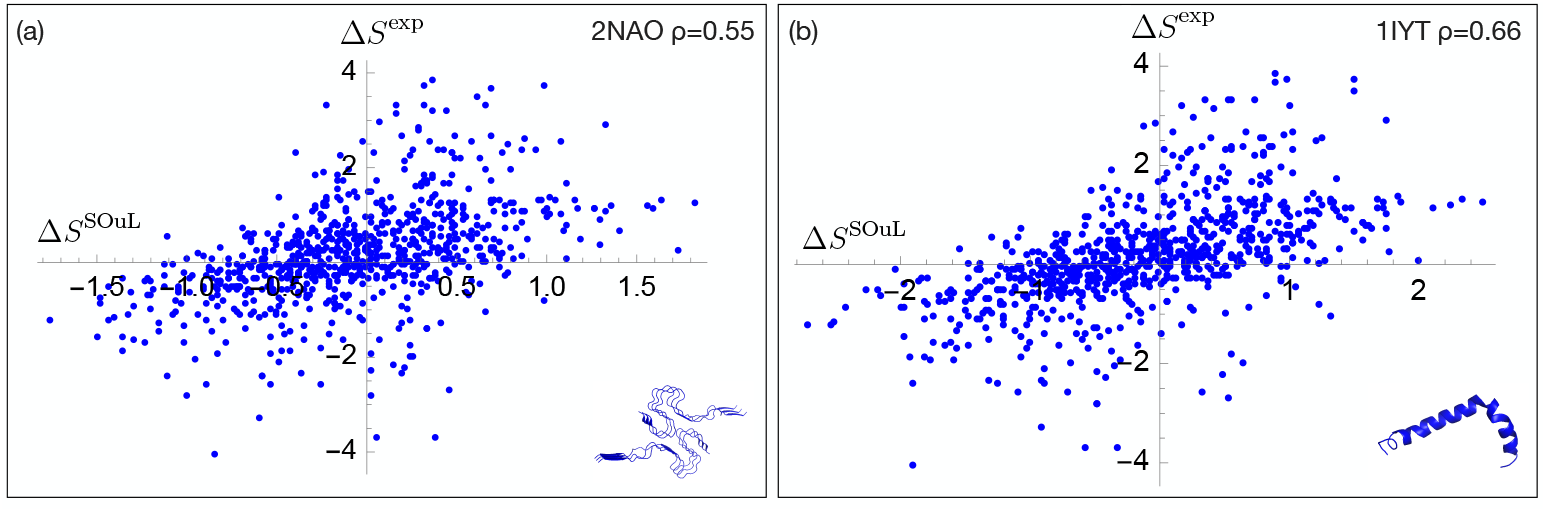
SOuLMuSiC performances on A*β*-42 variants using as input (a) the fibril structure PDB code 2NAO and (b) the monomeric protein in solution with PDB code 1IYT. The protein structures are shown on the bottom right.

As shown in the figure, the predicted Δ*S*^*SOuL*^ values are well correlated with the experimental Δ*S*^*exp*^ values, with Spearman correlation coefficients of *ρ* = 0.55 and 0.66 for the 2NAO and 1IYT structures, respectively. The *β* parameter in Eq. 6 is set to -1 for 2NAO and to +1 for 1IYT, as one represents the fibril structure and the other the protein in solution. Note that exchanging the *β* values results in a drastic drop in performance, with Spearman correlation decreasing to around 0.2. It is also interesting to note that the correlation between predictions obtained with the two structures that, despite having completely different conformations, is high with a Pearson coefficient of 0.84.

### Webserver

To make SOuLMuSiC accessible to the large academic community, we made it available through the webserver http://babylone.3bio.ulb.ac.be/SOuLMuSiC/. As SOuLMuSiC is structure-based, users must provide a 3D structure of the target protein as input. The user has the choice among three different methods for this: provide the PDB code [40] whose structure will be automatically retrieved from the PDB; the UniProt ID, in which case the corresponding structure will be retrieved from AlphaFold DB [80]; or upload his own structure in PDB format. Note that SOuLMuSiC takes into account all chains contained in the submitted structure file when computing the solubility score upon mutations, so users should provide only the chains of interest. The user also has to choose whether the provided structure is a globular protein or a insoluble macromolecular assembly.

Once the structure is submitted, the computation starts. SOuLMuSiC is very fast and performs the prediction of solubility for all single-site mutations in a protein in less than a minute. Upon completion of the calculation, a CSV file containing the results is sent to the email address provided upon submission. The result file contains, for all single-site mutations in all chains of the submitted protein structure, the solvent accessibility of the mutated residue and the Δ*S*^*SOuL*^ value.

## 4 Discussion

The optimization of protein solubility is one of the fundamental goals in any biotechnological process involving proteins. Despite the considerable efforts made over the last decades, it is still difficult to identify, whether experimentally or computationally, the impact of mutations on protein solubility. In this article, we have taken a step towards this goal by presenting SOuLMuSiC, our tool of the MuSiC suite [50, 64, 81, 82, 83] dedicated to predict the effect of single-site mutations on protein solubility. Our tool outperforms state-of-the-art approaches when tested on a highly curated dataset of mutations manually collected from the literature and has also been successfully validated on external datasets to assess the generalizability of our approach.

Additional aspects should be explored to further improve our approach. The development of accurate solubility predictors is hindered by the limited availability of mutational data on solubility. Expanding the dataset in the near future could facilitate the training of more complex models with a larger number of features and parameters. Deep mutagenesis data are certainly helpful, though their precise link to protein solubility is complex; solubility, while related to stability and aggregation, is a distinct property, and disentangling these factors is challenging, as we observed in the Results section. Another issue is related to data variability, since solubility measurements are performed using various experimental setups. Even though we made efforts to standardize all collected information in constructing our dataset, this variability remains a source of noise. Finally, the impact of environmental variables, such as pH or temperature, plays a significant role in protein solubility. Although not currently taken in account in SOuLMuSiC, incorporating these factors into the models could improve their performance.

Although SOuLMuSiC can still be optimized, its current version already achieves good performance. Given the significant challenges that solubility issues present in both academic and industrial processes, we are confident that SOuLMuSiC will be of great interest to the scientific community and will be used to rationally design new proteins with improved solubility, which, in turn, could help optimize a wide variety of biotechnological processes.

## Notes

### Competing Interest Statement

The authors have declared no competing interest.

## References

[1] Filipa Castro and Joana Ferreira. Advances in protein solubility and thermodynamics: quantification, instrumentation, and perspectives. CrystEngComm, 2023.

[2] Ryan M Kramer, Varad R Shende, Nicole Motl, C Nick Pace, and J Martin Scholtz. Toward a molecular understanding of protein solubility: increased negative surface charge correlates with increased solubility. Biophysical journal, 102(8):1907–1915, 2012.

[3] Mauno Vihinen. Solubility of proteins. ADMET and DMPK, 8(4):391–399, 2020.

[4] Steven J Shire, Zahra Shahrokh, and JUN Liu. Challenges in the development of high protein concentration formulations. Journal of pharmaceutical sciences, 93(6):1390–1402, 2004.

[5] Wim Jiskoot, Andrea Hawe, Tim Menzen, David B Volkin, and Daan JA Crommelin. Ongoing challenges to develop high concentration monoclonal antibody-based formulations for subcutaneous administration: Quo vadis? Journal of Pharmaceutical Sciences, 111(4):861–867, 2022.

[6] Agustín Correa and Pablo Oppezzo. Overcoming the solubility problem in e. coli: available approaches for recombinant protein production. Insoluble Proteins: Methods and Protocols, pages 27–44, 2015.

[7] Mahin Pouresmaeil and Shahnam Azizi-Dargahlou. Factors involved in heterologous expression of proteins in e. coli host. Archives of Microbiology, 205 (5):212, 2023.

[8] Christian Haass and Dennis J Selkoe. Soluble protein oligomers in neurodegeneration: lessons from the alzheimer’s amyloid β-peptide. Nature reviews Molecular cell biology, 8(2):101–112, 2007.

[9] Baraa A Hijaz and Laura A Volpicelli-Daley. Initiation and propagation of α-synuclein aggregation in the nervous system. Molecular neurodegeneration, 15:1–12, 2020.

[10] Venkata Pulla Rao Vendra, Ismail Khan, Sushil Chandani, Anbukkarasi Muniyandi, and Dorairajan Balasubramanian. Gamma crystallins of the human eye lens. Biochimica et Biophysica Acta (BBA)-General Subjects, 1860(1):333–343, 2016.

[11] Jennifer C Boatz, Matthew J Whitley, Mingyue Li, Angela M Gronenborn, and Patrick CA van der Wel. Cataract-associated p23t γd-crystallin retains a native-like fold in amorphous-looking aggregates formed at physiological ph. Nature Communications, 8(1):15137, 2017.

[12] Abhisek Mukherjee, Diego Morales-Scheihing, Peter C Butler, and Claudio Soto. Type 2 diabetes as a protein misfolding disease. Trends in molecular medicine, 21(7):439–449, 2015.

[13] Michel Goedert, Florence Clavaguera, and Markus Tolnay. The propagation of prion-like protein inclusions in neurodegenerative diseases. Trends in neurosciences, 33(7):317–325, 2010.

[14] Qingzhen Hou, Raphaël Bourgeas, Fabrizio Pucci, and Marianne Rooman. Computational analysis of the amino acid interactions that promote or decrease protein solubility. Scientific reports, 8(1):14661, 2018.

[15] Bikash K Bhandari, Paul P Gardner, and Chun Shen Lim. Solubility-weighted index: fast and accurate prediction of protein solubility. Bioinformatics, 36 (18):4691–4698, 2020.

[16] Jim Warwicker, Spyros Charonis, and Robin A Curtis. Lysine and arginine content of proteins: computational analysis suggests a new tool for solubility design. Molecular pharmaceutics, 11(1):294–303, 2014.

[17] Pedro Chan, Robin A Curtis, and Jim Warwicker. Soluble expression of proteins correlates with a lack of positively-charged surface. Scientific reports, 3 (1):3333, 2013.

[18] Dominic Esposito and Deb K Chatterjee. Enhancement of soluble protein expression through the use of fusion tags. Current opinion in biotechnology, 17 (4):353–358, 2006.

[19] Surinder Mohan Singh and Amulya Kumar Panda. Solubilization and refolding of bacterial inclusion body proteins. Journal of bioscience and bioengineering, 99(4):303–310, 2005.

[20] Luis Felipe Vallejo and Ursula Rinas. Strategies for the recovery of active proteins through refolding of bacterial inclusion body proteins. Microbial cell factories, 3:1–12, 2004.

[21] Anupam Singh, Vaibhav Upadhyay, Arun Kumar Upadhyay, Surinder Mohan Singh, and Amulya Kumar Panda. Protein recovery from inclusion bodies of escherichia coli using mild solubilization process. Microbial cell factories, 14: 1–10, 2015.

[22] Filipe SR Silva, Sara PO Santos, Roberto Meyer, Eduardo S Silva, Carina S Pinheiro, Neuza M Alcantara-Neves, and Luis GC Pacheco. In vivo cleavage of solubility tags as a tool to enhance the levels of soluble recombinant proteins in escherichia coli. Biotechnology and Bioengineering, 118(11):4159–4167, 2021.

[23] Tatsuya Niwa, Bei-Wen Ying, Katsuyo Saito, WenZhen Jin, Shoji Takada, Takuya Ueda, and Hideki Taguchi. Bimodal protein solubility distribution revealed by an aggregation analysis of the entire ensemble of escherichia coli proteins. Proceedings of the National Academy of Sciences, 106(11):4201–4206, 2009.

[24] Christophe N Magnan, Arlo Randall, and Pierre Baldi. Solpro: accurate sequence-based prediction of protein solubility. Bioinformatics, 25(17):2200–2207, 2009.

[25] Federico Agostini, Davide Cirillo, Carmen Maria Livi, Riccardo Delli Ponti, and Gian Gaetano Tartaglia. cc sol omics: A webserver for solubility prediction of endogenous and heterologous expression in escherichia coli. Bioinformatics, 30(20):2975–2977, 2014.

[26] Max Hebditch, M Alejandro Carballo-Amador, Spyros Charonis, Robin Curtis, and Jim Warwicker. Protein–sol: a web tool for predicting protein solubility from sequence. Bioinformatics, 33(19):3098–3100, 2017.

[27] Qingzhen Hou, Jean Marc Kwasigroch, Marianne Rooman, and Fabrizio Pucci. Solart: a structure-based method to predict protein solubility and aggregation. Bioinformatics, 36(5):1445–1452, 2020.

[28] Jiri Hon, Martin Marusiak, Tomas Martinek, Antonin Kunka, Jaroslav Zen-dulka, David Bednar, and Jiri Damborsky. Soluprot: prediction of soluble protein expression in escherichia coli. Bioinformatics, 37(1):23–28, 2021.

[29] Daniele Raimondi, Gabriele Orlando, Piero Fariselli, and Yves Moreau. Insight into the protein solubility driving forces with neural attention. PLoS computational biology, 16(4):e1007722, 2020.

[30] Vineet Thumuluri, Hannah-Marie Martiny, Jose J Almagro Armenteros, Jesper Salomon, Henrik Nielsen, and Alexander Rosenberg Johansen. Netsolp: predicting protein solubility in escherichia coli using language models. Bioinformatics, 38(4):941–946, 2022.

[31] Jan Velecky, Marie Hamsikova, Jan Stourac, Milos Musil, Jiri Damborsky, David Bednar, and Stanislav Mazurenko. Soluprotmutdb: A manually curated database of protein solubility changes upon mutations. Computational and Structural Biotechnology Journal, 20:6339–6347, 2022.

[32] Pietro Sormanni, Francesco A Aprile, and Michele Vendruscolo. The camsol method of rational design of protein mutants with enhanced solubility. Journal of molecular biology, 427(2):478–490, 2015.

[33] Lisanna Paladin, Damiano Piovesan, and Silvio CE Tosatto. Soda: prediction of protein solubility from disorder and aggregation propensity. Nucleic acids research, 45(W1):W236–W240, 2017.

[34] Yang Yang, Lianjie Zeng, and Mauno Vihinen. Pon-sol2: Prediction of effects of variants on protein solubility. International Journal of Molecular Sciences, 22(15):8027, 2021.

[35] Joana Ferreira and Filipa Castro. Advances in protein solubility and thermodynamics: quantification, instrumentation, and perspectives. CrystEngComm, 25(46):6388–6404, 2023.

[36] Saul R Trevino, J Martin Scholtz, and C Nick Pace. Measuring and increasing protein solubility. Journal of pharmaceutical sciences, 97(10):4155–4166, 2008.

[37] Marc Oeller, Ryan Kang, Rosie Bell, Hannes Ausserwoger, Pietro Sormanni, and Michele Vendruscolo. Sequence-based prediction of ph-dependent protein solubility using camsol. Briefings in Bioinformatics, 24(2):bbad004, 2023.

[38] BA Chrunyk, J Evans, J Lillquist, P Young, and R Wetzel. Inclusion body for-mation and protein stability in sequence variants of interleukin-1 beta. Journal of Biological Chemistry, 268(24):18053–18061, 1993.

[39] Amir Y Mirarefi and Charles F Zukoski. Gradient diffusion and protein solubility: use of dynamic light scattering to localize crystallization conditions. Journal of crystal growth, 265(1-2):274–283, 2004.

[40] Helen M Berman, John Westbrook, Zukang Feng, Gary Gilliland, Talapady N Bhat, Helge Weissig, Ilya N Shindyalov, and Philip E Bourne. The protein data bank. Nucleic acids research, 28(1):235–242, 2000.

[41] John Jumper, Richard Evans, Alexander Pritzel, Tim Green, Michael Figurnov, Olaf Ronneberger, Kathryn Tunyasuvunakool, Russ Bates, Augustin Zídek, Anna Potapenko, et al. Highly accurate protein structure prediction with alphafold. Nature, 596(7873):583–589, 2021.

[42] Andrew Waterhouse, Martino Bertoni, Stefan Bienert, Gabriel Studer, Gerardo Tauriello, Rafal Gumienny, Florian T Heer, Tjaart A P de Beer, Christine Rempfer, Lorenza Bordoli, et al. Swiss-model: homology modelling of protein structures and complexes. Nucleic acids research, 46(W1):W296–W303, 2018.

[43] Fabrizio Pucci, Katrien V Bernaerts, Jean Marc Kwasigroch, and Marianne Rooman. Quantification of biases in predictions of protein stability changes upon mutations. Bioinformatics, 34(21):3659–3665, 2018.

[44] András Fiser and Andrej Šali. Modeller: generation and refinement of homology-based protein structure models. In Methods in enzymology, volume 374, pages 461–491. Elsevier, 2003.

[45] Christina Rother, Alexander Gutmann, Ramakrishna Gudiminchi, Hansjörg Weber, Alexander Lepak, and Bernd Nidetzky. Biochemical characterization and mechanistic analysis of the levoglucosan kinase from lipomyces starkeyi. ChemBioChem, 19(6):596–603, 2018.

[46] Justin R Klesmith, John-Paul Bacik, Emily E Wrenbeck, Ryszard Michalczyk, and Timothy A Whitehead. Trade-offs between enzyme fitness and solubility illuminated by deep mutational scanning. Proceedings of the National Academy of Sciences, 114(9):2265–2270, 2017.

[47] Mireia Seuma, Ben Lehner, and Benedetta Bolognesi. An atlas of amyloid aggregation: the impact of substitutions, insertions, deletions and truncations on amyloid beta fibril nucleation. Nature Communications, 13(1):7084, 2022.

[48] Sanzo Miyazawa and Robert L Jernigan. Residue–residue potentials with a favorable contact pair term and an unfavorable high packing density term, for simulation and threading. Journal of molecular biology, 256(3):623–644, 1996.

[49] Yves Dehouck, Dimitri Gilis, and Marianne Rooman. A new generation of statistical potentials for proteins. Biophysical journal, 90(11):4010–4017, 2006.

[50] Yves Dehouck, Jean Marc Kwasigroch, Dimitri Gilis, and Marianne Rooman. Popmusic 2.1: a web server for the estimation of protein stability changes upon mutation and sequence optimality. BMC bioinformatics, 12:1–12, 2011.

[51] Qingzhen Hou, Fabrizio Pucci, François Ancien, Jean Marc Kwasigroch, Raphaël Bourgeas, and Marianne Rooman. Swotein: a structure-based approach to predict stability strengths and weaknesses of proteins. Bioinformatics, 37(14):1963–1971, 2021.

[52] Georgios A Dalkas, Fabian Teheux, Jean Marc Kwasigroch, and Marianne Rooman. Cation–π, amino–π, π–π, and h-bond interactions stabilize antigen– antibody interfaces. Proteins: Structure, Function, and Bioinformatics, 82(9): 1734–1746, 2014.

[53] Marianne J Rooman, Jean-Pierre A Kocher, and Shoshana J Wodak. Prediction of protein backbone conformation based on seven structure assignments: influence of local interactions. Journal of molecular biology, 221(3):961–979, 1991.

[54] Jean-Pierre A Kocher, Marianne J Rooman, and Shoshana J Wodak. Factors influencing the ability of knowledge-based potentials to identify native sequence-structure matches. Journal of molecular biology, 235(5):1598–1613, 1994.

[55] William C Wimley and Stephen H White. Experimentally determined hydrophobicity scale for proteins at membrane interfaces. Nature structural biology, 3(10):842–848, 1996.

[56] Jack Kyte and Russell F Doolittle. A simple method for displaying the hydropathic character of a protein. Journal of molecular biology, 157(1):105–132, 1982.

[57] JOEL Janin. Surface and inside volumes in globular proteins. Nature, 277 (5696):491–492, 1979.

[58] Charles Tanford. Physical chemistry of macromolecules. John Wiley & Sons, Incorporated, 1966.

[59] Kevin L Shaw, Gerald R Grimsley, Gennady I Yakovlev, Alexander A Makarov, and C Nick Pace. The effect of net charge on the solubility, activity, and stability of ribonuclease sa. Protein Science, 10(6):1206–1215, 2001.

[60] Kuo-Chen Chou. Using amphiphilic pseudo amino acid composition to predict enzyme subfamily classes. Bioinformatics, 21(1):10–19, 2005.

[61] Nan Xiao, Dong-Sheng Cao, Min-Feng Zhu, and Qing-Song Xu. protr/protrweb: R package and web server for generating various numerical representation schemes of protein sequences. Bioinformatics, 31(11):1857–1859, 2015.

[62] J Meier, R Rao, R Verkuil, J Liu, T Sercu, and A Rives. Language models enable zero-shot prediction of the effects of mutations on protein function. biorxiv, 2021.07.09.450648, 2021.

[63] Baris E Suzek, Hongzhan Huang, Peter McGarvey, Raja Mazumder, and Cathy H Wu. Uniref: comprehensive and non-redundant uniprot reference clusters. Bioinformatics, 23(10):1282–1288, 2007.

[64] Fabrizio Pucci, Raphaël Bourgeas, and Marianne Rooman. Predicting protein thermal stability changes upon point mutations using statistical potentials: Introducing hotmusic. Scientific reports, 6(1):23257, 2016.

[65] Dimitri Gilis and Marianne Rooman. Predicting protein stability changes upon mutation using database-derived potentials: solvent accessibility determines the importance of local versus non-local interactions along the sequence. Journal of molecular biology, 272(2):276–290, 1997.

[66] Wolfram Research, Inc. Mathematica, Version 14.0. URL https://www.wolfram.com/mathematica. Champaign, IL, 2024.

[67] Dinara R Usmanova, Natalya S Bogatyreva, Joan Ariño Bernad, Aleksandra A Eremina, Anastasiya A Gorshkova, German M Kanevskiy, Lyubov R Lonishin, Alexander V Meister, Alisa G Yakupova, Fyodor A Kondrashov, et al. Self-consistency test reveals systematic bias in programs for prediction change of stability upon mutation. Bioinformatics, 34(21):3653–3658, 2018.

[68] Matsvei Tsishyn, Fabrizio Pucci, and Marianne Rooman. Quantification of biases in predictions of protein–protein binding affinity changes upon mutations. Briefings in bioinformatics, 25(1):bbad491, 2024.

[69] Lavi S Bigman and Yaakov Levy. Proteins: molecules defined by their tradeoffs. Current opinion in structural biology, 60:50–56, 2020.

[70] Aron Broom, Zachary Jacobi, Kyle Trainor, and Elizabeth M Meiering. Computational tools help improve protein stability but with a solubility tradeoff. Journal of Biological Chemistry, 292(35):14349–14361, 2017.

[71] Pauline Hermans, Matsvei Tsishyn, Martin Schwersensky, Marianne Rooman, and Fabrizio Pucci. Exploring evolution to uncover insights into protein mutational stability. Molecular Biology and Evolution, page msae267, 2024.

[72] Qingzhen Hou, Marianne Rooman, and Fabrizio Pucci. Enzyme stability-activity trade-off: new insights from protein stability weaknesses and evolutionary conservation. Journal of chemical theory and computation, 19(12): 3664–3671, 2023.

[73] Fabrizio Chiti and Christopher M Dobson. Protein misfolding, functional amyloid, and human disease. Annu. Rev. Biochem., 75(1):333–366, 2006.

[74] Nikolaos Louros, Joost Schymkowitz, and Frederic Rousseau. Mechanisms and pathology of protein misfolding and aggregation. Nature Reviews Molecular Cell Biology, 24(12):912–933, 2023.

[75] Federico Agostini, Michele Vendruscolo, and Gian Gaetano Tartaglia. Sequence-based prediction of protein solubility. Journal of molecular biology, 421(2-3):237–241, 2012.

[76] H Jane Dyson and Peter E Wright. Intrinsically unstructured proteins and their functions. Nature reviews Molecular cell biology, 6(3):197–208, 2005.

[77] Marielle Aulikki Walti, Francesco Ravotti, Hiromi Arai, Charles G Glabe, Joseph S Wall, Anja Bockmann, Peter Guntert, Beat H Meier, and Roland Riek. Atomic-resolution structure of a disease-relevant aβ (1–42) amyloid fibril. Proceedings of the National Academy of Sciences, 113(34):E4976–E4984, 2016.

[78] Orlando Crescenzi, Simona Tomaselli, Remo Guerrini, Severo Salvadori, Anna M D’Ursi, Piero Andrea Temussi, and Delia Picone. Solution structure of the alzheimer amyloid β-peptide (1–42) in an apolar microenvironment: Similarity with a virus fusion domain. European journal of biochemistry, 269(22): 5642–5648, 2002.

[79] Vanessa E Gray, Katherine Sitko, Floriane Z Ngako Kameni, Miriam Williamson, Jason J Stephany, Nicholas Hasle, and Douglas M Fowler. Elucidating the molecular determinants of aβ aggregation with deep mutational scanning. G3: Genes, Genomes, Genetics, 9(11):3683–3689, 2019.

[80] Mihaly Varadi, Damian Bertoni, Paulyna Magana, Urmila Paramval, Ivanna Pidruchna, Malarvizhi Radhakrishnan, Maxim Tsenkov, Sreenath Nair, Milot Mirdita, Jingi Yeo, et al. Alphafold protein structure database in 2024: providing structure coverage for over 214 million protein sequences. Nucleic acids research, 52(D1):D368–D375, 2024.

[81] Yves Dehouck, Jean Marc Kwasigroch, Marianne Rooman, and Dimitri Gilis. Beatmusic: prediction of changes in protein–protein binding affinity on mutations. Nucleic acids research, 41(W1):W333–W339, 2013.

[82] François Ancien, Fabrizio Pucci, Maxime Godfroid, and Marianne Rooman. Prediction and interpretation of deleterious coding variants in terms of protein structural stability. Scientific reports, 8(1):4480, 2018.

[83] Matsvei Tsishyn, Gabriel Cia, Pauline Hermans, Jean Kwasigroch, Marianne Rooman, and Fabrizio Pucci. Fitmusic: leveraging structural and (co) evolutionary data for protein fitness prediction. Human genomics, 18(1):36, 2024.

